# Deterministic response strategies in trial-and-error learning

**DOI:** 10.1101/258459

**Authors:** Holger Mohr, Katharina Zwosta, Dimitrije Markovic, Sebastian Bitzer, Uta Wolfensteller, Hannes Ruge

## Abstract

Trial-and-error learning is a universal strategy for establishing which actions are beneficial or harmful in new environments. However, learning stimulus-response associations solely via trial-and-error is often suboptimal, as in many settings dependencies among stimuli and responses can be exploited to increase learning efficiency. Previous studies have shown that in settings featuring such dependencies, humans typically engage high-level cognitive processes and employ advanced learning strategies to improve their learning efficiency. Here we analyze in detail the initial learning phase of a sample of human subjects (N = 85) performing a trial-and-error learning task with deterministic feedback and hidden stimulus-response dependencies. Using computational modeling, we find that the standard Q-learning model cannot sufficiently explain human learning strategies in this setting. Instead, newly introduced deterministic response models, which are theoretically optimal and transform stimulus sequences unambiguously into response sequences, provide the best explanation for 50.6% of the subjects. Most of the remaining subjects either show a tendency towards generic optimal learning (21.2%) or at least partially exploit stimulus-response dependencies (22.3%), while a few subjects (5.9%) show no clear preference for any of the employed models. After the initial learning phase, asymptotic learning performance during the subsequent practice phase is best explained by the standard Q-learning model. Our results show that human learning strategies in trial-and-error learning go beyond merely associating stimuli and responses via incremental reinforcement. Specifically during initial learning, high-level cognitive processes support sophisticated learning strategies that increase learning efficiency while keeping memory demands and computational efforts bounded. The good asymptotic fit of the Q-learning model indicates that these cognitive processes are successively replaced by the formation of stimulus-response associations over the course of learning.

## Introduction

Learning rewarded stimulus-response associations via trial-and-error can be a powerful strategy, which has been employed successfully in complex learning tasks (1). However, human learning strategies in trial-and-error learning tasks typically go beyond merely associating stimuli and responses via reinforcement. Instead, it has been shown that humans employ high-level cognitive capabilities like working memory and attention to make learning more efficient by exploiting hidden or overt structure in the environment (2–6). For example, it was shown that subjects can quickly reactivate previously learned response strategies (7) and incorporate information on unselected response options to improve learning efficiency (8–10). Building on a long history of research on associative learning (11, 12), recent studies increasingly employed advanced modeling approaches like reinforcement learning or Bayesian and Hidden Markov models to explain human learning strategies in various learning tasks (13–15). Specifically, Q-learning models have been adapted or extended to account for high-level cognitive processes engaged during learning. For instance, Collins et al. have shown in a series of studies that by adding a working memory module to the standard Q-learning model, human learning can be better explained than by pure associative learning (2, 16, 17), see also (18). Leong et al. showed that an extended reinforcement learning model with separate weights for different stimulus dimensions can capture attention-related processes in a trial-and-error learning task (19). Moreover, several studies have shown that humans incorporate implicit relations and hidden task structure into their learning strategy to make learning more efficient (4, 20, 21). Specifically in probabilistic settings, it was shown that when updating internal beliefs about reward probabilities, humans integrate information about unchosen stimuli-response pairs into the updating process both in tasks overtly presenting the outcome of the unchosen options and in tasks with implicit outcome contingencies (8–10, 14, 15, 22–24). Using probabilistic reward schemes, including fluctuating reward probabilities or dependencies, these studies showed that modified Q-learning, Bayesian or Hidden Markov models, approximating optimal performance in the respective learning tasks, outperformed the standard Q-learning model serving as a baseline for comparison with the more sophisticated models.

Here we show that even in a simple learning task with deterministic feedback, human learning strategies can be surprisingly complex. Specifically, we introduce novel deterministic response pattern models to test whether subjects explore response options in a fix order during the initial learning phase. These deterministic models are compared to three alternative models, which are the standard Q-learning model reflecting pure associative learning, a generic optimal model that fully exploits hidden stimulus-response dependencies, and an intermediate model that exploits dependencies less efficiently than the optimal learning model but more efficiently than the Q-learning model.

## Results

Subjects performed a simple stimulus-response learning task with deterministic feedback (N = 85), see also (25). All subjects were informed about the purpose and procedure of the experiment and gave written informed consent prior to taking part in the experiment, in accordance with the Declaration of Helsinki. In each learning block, a novel set of four stimuli was introduced and subjects had to learn the correct responses to the four stimuli (see Figure 1). The set of responses remained constant across blocks and consisted of the four keys *d,f,k,l* on a computer keyboard, corresponding to the left middle, left index, right index and right middle finger. Each stimulus was associated with a unique correct response, i.e. stimuli mapped onto responses one-to-one. Feedback was given deterministically, i.e. correct/incorrect responses were invariably indicated by positive/negative feedback.

**Fig. 1.**
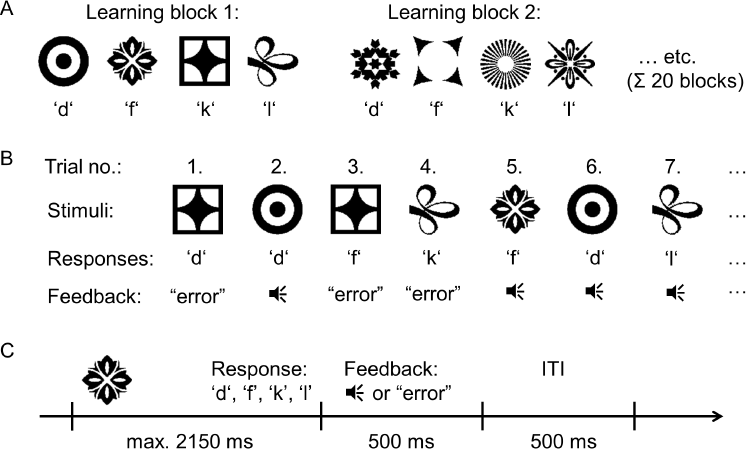
The trial-and-error learning task. **A:** In each learning block, subjects had to learn the correct responses to four novel stimuli (N = 85). Stimuli mapped onto responses one-to-one, i.e. each stimulus was associated with a unique correct response. Each subject performed 20 learning blocks. **B:** Stimuli were presented in randomized order, and subjects responded with one of the four keys ***d, f, k, l*** on a computer keyboard. After response selection, subjects were provided with feedback indicating a correct response via auditory feedback or an incorrect response via the word ‘error’ written on the screen. Learning blocks ended when each stimulus had been performed correctly eight times or maximally after 70 trials. **C:** Response times were limited to 2150 ms, feedback was presented for 500 ms, followed by an inter-trial interval of 500 ms.

**Q-learning.** The standard Q-learning model served as a baseline for comparison with more sophisticated models (26). In Q-learning, associations between stimuli and responses are expressed as Q-values (action values or associative weights), which were set to zero initially and were updated after each trial with learning rate *α* ∈ (0,1] based on the following learning rule:

After positive feedback for stimulus-response pair *S_i_, R_j_*:

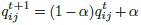

After negative feedback for stimulus-response pair *S_i_*, *R_j_*:

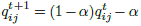

Response probabilities were determined via the softmax response selection rule with noise parameter *τ* ≥ 0:

Given *S_i_*, the probability for selecting response *Rj* was:

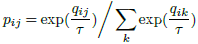

For the special case *τ* = 0 (noise-free response selection), responses were selected uniformly among the responses with maximal Q-values.

Note that the Q-learning model updates its associative weights for each stimulus-response (S-R) pair separately, i.e. independently of the other stimulus-response pairs. Hence, this model cannot directly capture dependencies among different stimulus-response-outcome (S-R-O) combinations. Specifically, Q-learning cannot exploit the one-to-one property of the S-R mappings, i.e. the fact that once a response has been associated with a stimulus, this response can be excluded for the other three stimuli.

**Free optimal play (FOP).** Based on the literature discussed in the introduction, we hypothesized that subjects may show a tendency towards optimal behavior, i.e. exploit the dependencies among S-R pairs, rather than learning S-R associations independently via reinforcement. In order to maximize expected reward while concurrently minimizing expected uncertainty, the following optimal learning strategy can be employed: Given the 4 stimuli and 4 responses, there are 4*3*2 = 24 possible S-R mappings. At the beginning of a learning block, there is no evidence against any of these 24 mappings, thus the probability for each mapping is assumed to be 1/24 (see Figure 2). After each trial, the set of S-R mappings that are consistent with the observed S-R-O history is updated. For each S-R pair, the probability of being correct can be computed by averaging across the set of consistent SR mappings. Selecting the most likely responses according to this procedure maximizes expected reward and minimizes expected uncertainty, see Appendix for a technical discussion.

**Fig. 2.**
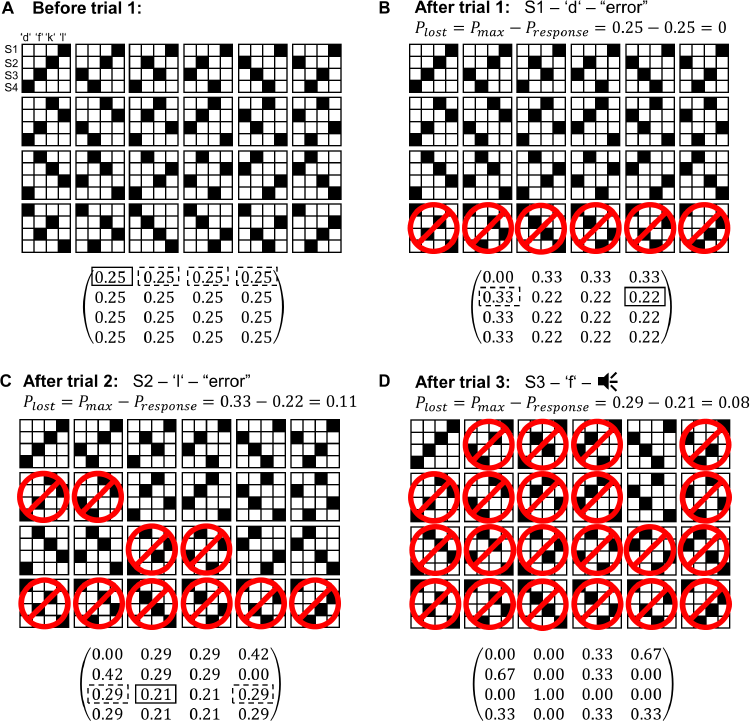
Computation of response probabilities. The four stimuli can map one-to-one onto the four responses in 24 different ways, depicted by the 24 matrices, with rows corresponding to stimuli and columns to responses. As feedback was deterministic, reward was delivered either with probability zero or one, indicated by the white and black squares, respectively. Overall probabilities (shown below the binary matrices) could be computed by averaging across the mappings that were consistent with the S-R-O history. **A:** At the beginning of a learning block, all 24 mappings were included in the set of consistent mappings. In the presented example, the subject chose response ***d*** in the first trial (solid box), which is optimal (i.e. provides the maximal likelihood of being rewarded), as were the other three response options (dashed boxes). **B:** The resulting negative feedback led to the exclusion of all S-R mappings that mapped stimulus S1 onto response ***d*** (indicated by the red no sign). In the next trial, the subject responded to stimulus S2 with l, resulting in negative feedback again. This response was not optimal, as response d was more likely than response l. **C:** Based on the feedback information, four additional mappings could be excluded. In the third trial, the subject responded with ***f*** to stimulus S3, which was correct (but not optimal a-priori). **D:** Only three S-R mappings are consistent with the S-R-O history at this point. Eventually, only the correct S-R mapping will remain. See Appendix for a technical discussion of the procedure.

The strategy of selecting a response that is maximally likely to be correct is termed free optimal play (FOP) in the following. Note that several responses can be maximally likely, i.e. this learning strategy does not necessarily determine a unique response. As this procedure required tracking the consistency of all 24 S-R mappings and computing averages across subsets of S-R mappings, it seemed unlikely that the subjects implemented this strategy. Yet, we hypothesized that there might be a trend towards this optimal strategy. Indeed, if subjects occasionally exploited the one-to-one property of the SR mappings, free optimal play might provide a better fit to the data than Q-learning.

As in Q-learning, response selection probabilities in the FOP model were determined by a softmax rule:

Given *S_i_*, the probability for selecting response *R_j_* was:

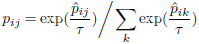

with 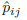 denoting the probabilities as computed by the FOP scheme (Figure 2).

**Binarized play (BP).** To test whether the subjects tracked the fine-grained differences between response probabilities as provided by FOP, or alternatively, only excluded responses that had already been assigned to a different stimulus, we implemented a simpler version of free optimal play, termed binarized play (BP), that was no longer optimal. The probabilities 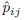 as computed by the FOP model were transformed into a simplified distribution by making all nonzero probabilities uniform, i.e. for a given stimulus *S*_*i*_, the BP probabilities 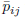 were defined as

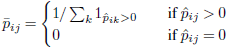

For example, for a given stimulus *S_i_*, a vector of response probabilities 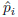 = (0.6,0,0.3,0.1), computed according to FOP, was transformed into 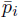 = (0.33,0,0.33,0.33). Response selection probabilities were again computed via the softmax rule:

Given *S_i_*, the probability for selecting response *R_j_* was:

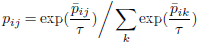

**Deterministic response patterns (DRPs).** Instead of tracking all 24 S-R mappings as required by FOP, the task could also be optimally performed with reduced memory and computational demands by means of deterministic response strategies. In contrast to FOP, responses are tested in a fixed order for all stimuli, for instance by going from left to right on the keyboard (*dfkl*). In case of negative feedback, the next response according to the response order is tested at the subsequent presentation of the stimulus. Alternatively, if the response is correct, it is logged in for the respective stimulus, and the response is excluded for the remaining stimuli, i.e. the next response to test in the fixed order is the next response that has not yet been assigned to any stimulus (see Figure 3). From a theoretical point of view, the order by which the responses are tested is arbitrary, i.e. any of the 24 possible response orders could be used to perform the task. However, we hypothesized that from a human perspective, certain response orders, like *dfkl* (going from left to right) or *lkfd* (going from right to left), might be easier to implement than others.

**Fig. 3.**
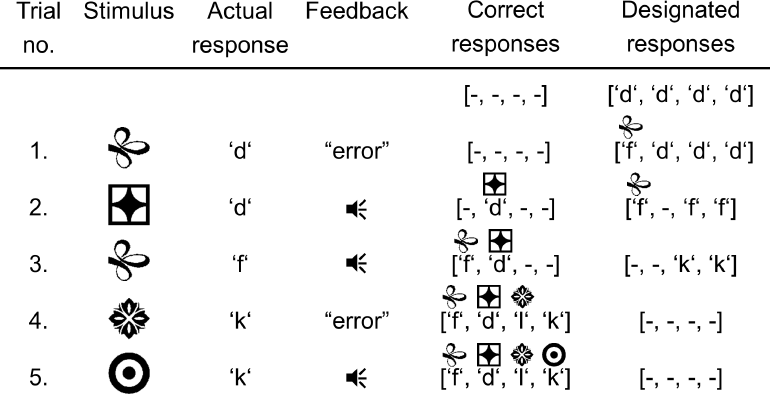
Example for a deterministic response pattern. Using the response order dfkl, the deterministic response pattern unfolds in the following way: Before trial 1, all four stimuli are to be responded by the first response of the response order, which is response d in this example. Due to the negative feedback in trial 1, the designated response for stimulus S1 is set to the next response according to the response order, which is f in this example. In the second trial, responding with d to stimulus S2 results in positive feedback, thus response d is logged in as the correct response for this stimulus. Importantly, due to the one-to-one property of the S-R mappings, the response d is blocked for the other three stimuli, thus the designated responses for stimuli S3 and S4 are set to the next unoccupied response, which is f. In trial 3, response f is logged in for stimulus S1, and the designated responses for stimuli S3, S4 are set to the next response according to the response order, which is response k in this example. In trial 4, responding with k to stimulus S3 results in negative feedback. At this point, again due to the one-to-one property of the S-R mapping, one can conclude that the correct response to stimulus S3 must be l. Moreover, although stimulus S4 has not yet been presented at this point, its correct response k can already be inferred.

The deterministic response pattern (DRP) models were implemented as follows: For a given stimulus *S_i_*, the response *R_j_* determined by the respective response order (either the designated or correct response) was set to probability one (i.e. 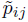 = 1) and the other three responses were set to probability zero (i.e. 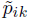 = 0 for *K* ≠ *j*). This degenerate distribution was transformed into a response selection probability distribution via the softmax rule:

Given *S_i_*, the probability for selecting response *R_j_* was:

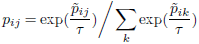

Under the presence of response selection noise (*τ* > 0), the updating procedure was defined in the following way: If the selected response deviated from the designated response due to response selection noise, only positive feedback led to an update, whereas negative feedback left the internal state of the model unchanged. This procedure was motivated by the fact that a deviation from the DRP could either occur as a backward deviation with respect to the response order, in which case the deviating response had been falsified before and updating was not necessary, or as a forward deviation, and in this case an update based on negative feedback would involve an invalid jump over possibly correct responses, thereby potentially corrupting the DRP procedure.

**Maximum likelihood estimates.** All models were fitted to the data by maximizing the log-likelihood of the data given the models, i.e. parameters were selected such that the actual responses were maximally likely given the models. The models were fitted on data of the initial learning phase of the learning blocks 6 to 20 (i.e. excluding blocks 1 to 5) to ensure that learning strategies had stabilized, since subjects had not been instructed on the one-to-one property of the S-R mappings before performing the task and thus had to adapt their learning strategies within the first few blocks. Models were fitted on data of the initial learning phase, which started at trial 1 and ended when all four stimuli were performed correctly at least once, i.e. the trial in which the fourth stimulus was performed correctly for the first time marked the end of the initial learning phase in each learning block. This was motivated by the fact that the DRP, FOP and BP models make no specific predictions for the subsequent practice phase following initial learning, besides the general prediction that correct responses are selected, up to a certain degree of fidelity determined by the noise parameter *τ*. Note that in contrast to the other models, the Q-learning model does make specific predictions for the practice phase, since S-R association strengths continuously increase with every correct repetition during the practice phase. Model parameters, consisting of the response selection noise *τ* ∈ (0,1/6.0,1/5.8,1/5.6,…, 1/0.2) for the DRP, FOP, BP and Q-learning models, and the learning rate *α* ∈ (0.05,0.10,0.15,*…*, 1.00) for the Q-learning model, were fitted separately for each subject on the initial learning phases of the learning blocks 6 to 20.

**Model comparisons.** Based on the maximum likelihood estimates, we determined for each subject which model provided the best fit to the initial learning phase, i.e. which model obtained the highest log-likelihood score (see Supplementary Figure S1). As expected, the response orders *dfkl* (going from left to right on the keyboard) and *lkfd* (from right to left) were ranked first and second among the DRP models, while the third-ranked response order was *kfdl*, which corresponds to the rather implausible sequence right index finger, left index finger, left middle finger, right middle finger, indicating a false positive hit for this response pattern. Thus, to be on the conservative side and avoid excessive statistical testing, we discarded all response orders but *dfkl* and *lkfd* and constrained our model space to the five models DRP *dfkl*, DRP *lkfd*, FOP, BP and Q-learning for subsequent analyses.

Only reporting which model scored the highest likelihood is in general not very informative, since the difference between the log-likelihood scores of the best and second best model can be arbitrarily small. Thus, we tested subject-wise whether one of the five models fitted the initial learning phase significantly better than competing models by conducting nonparametric Wilcoxon signed-rank tests across the log-likelihood values of the 15 learning blocks of interest. The five models were compared in an order corresponding to the quality of their predictions: As the DRP models make the most specific predictions on the learning strategy, we first tested for each subject whether either the DRP *dfkl* or DRP *lkfd* model fitted significantly better than the respective competing four models by conducting four pairwise one-sided Wilcoxon signed-rank tests across the 15 learning blocks of interest, using a significance threshold of P < 0.05. That is, if the DRP *dfkl* model fitted the initial learning phase of a given subject significantly better than the DRP lkfd, FOP, BP and Q-learning models (all four tests resulted in P < 0.05), the respective subject was assigned to the DRP subsample. Subjects were also assigned to the DRP subsample if the DRP lkfd model fitted significantly better than the DRP dfkl, FOP, BP and Q-learning models. If neither the DRP *dfkl* nor the DRP *lkfd* model outperformed the competing models, we tested whether the FOP model fitted significantly better than the BP and Q-learning models, as the FOP model makes more specific predictions than BP and Q-learning. That is, subjects were assigned to the FOP subsample if the FOP model fitted the initial learning phase significantly better than the BP and Q-learning models (both tests resulted in P < 0.05). For the remaining subjects, we tested whether the BP model fitted significantly better than Q-learning on the initial learning phase, and subjects were assigned to the BP subsample if this was the case. Finally, we tested whether Q-learning fitted significantly better than FOP or BP on the remaining subjects, and those subjects were assigned to the Q-learning subsample.

Based on this procedure, we found that the DRP models provided the best fit for 43 of 85 subjects (50.6%), with 36 subjects following the *dfkl* pattern and 7 subjects following the *lkfd* pattern (see Figure 4). A tendency towards generic optimal learning, as expressed by a better fit of FOP than BP and Q-learning, was found for 18 subjects (21.2%), while 19 subjects (22.3%) exploited the stimulus-response dependencies at least partially, as indicated by a better fit of BP than Q-learning. The remaining 5 subjects (5.9%) were assigned to none of the model-specific subsamples. Specifically, Q-learning did not fit significantly better than FOP or BP on the initial learning phase for any subject. Model parameter estimates are shown, separately for the three subsamples, in Supplementary Figure S2.

**Fig. 4.**
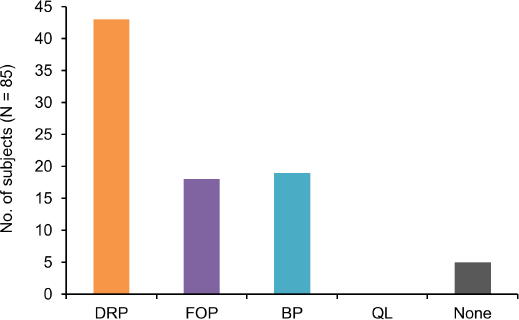
Result of the model comparison procedure. The trial-and-error learning task was performed by N = 85 subjects. For each subject, it was tested in descending order (see main text for details) which model provided the best fit for the initial learning phase. For 43 subjects (50.6%), the DRP models outperformed the FOP, BP and Q-learning models, with 36 subjects following the ***dfkl*** response pattern and 7 subjects following the ***lkfd*** pattern. Of the remaining subjects, 18 subjects (21.2%) showed a tendency towards generic optimal learning, while 19 subjects (22.3%) partially exploited stimulus-response dependencies. Q-learning was never significantly better than FOP or BP on the initial learning phase. Five subjects (5.9%) could not be assigned to a model-specific subsample.

**Learning curves.** Besides predictiveness, the generative performance of computational models is an important indicator for their ability to explain effects observed in the actual data (27). To evaluate the generative performance of the five models, we generated response data with N = 1000 repetitions for each learning block, using the respective subject-specific maximum likelihood model parameters. To evaluate the generative performance of the models in terms of learning dynamics, we compared the learning curves generated by the models with the actual learning curves of the subjects (see Figure 5). While the FOP and BP models provided better fits than Q-learning for the initial learning phase on all three sub-samples, the DRP models further improved the fit compared to FOP and BP within the first few trials on the DRP subsample. The Q-learning model provided the best asymptotic fit on all three subsamples, as the DRP, FOP and BP models made no specific predictions for the practice phase following initial learning beyond the general prediction that correct responses are selected with a certain degree of fidelity determined by the response selection noise parameter *τ*.

**Fig. 5.**
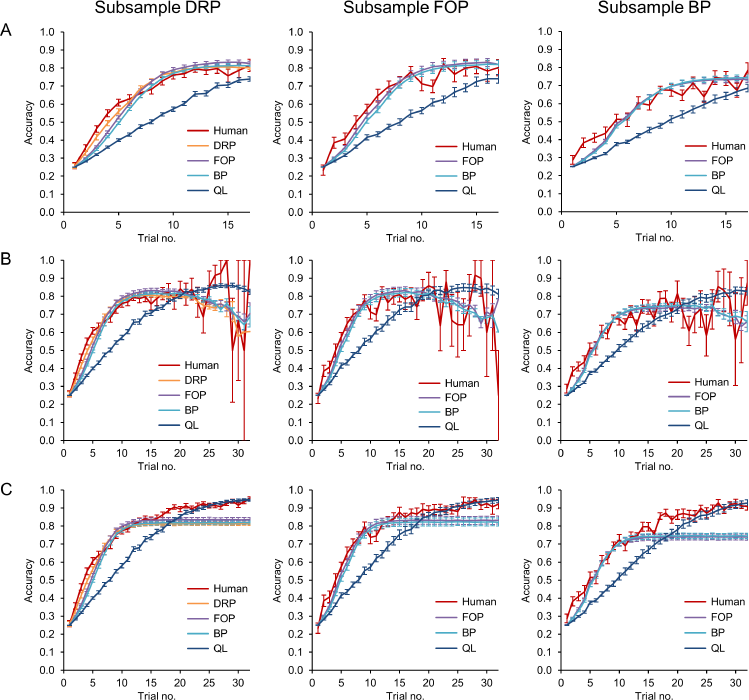
Learning curves for the three subsamples. **A:** Learning curves of the initial learning phase from trial 1 to 17. For the DRP subsample, the DRP, FOP and BP models provided a markedly better fit to the human learning curve than the Q-learning model. The DRP models improved the fit compared to the FOP and BP models for the first few trials. Within the FOP subsample, again both FOP and BP outperformed Q-learning, with the FOP model providing a marginally better fit than the BP model. For the BP subsample, the FOP and BP learning curves were indistinguishable but again fitted markedly better than Q-learning. Vertical lines indicate standard errors of the mean. **B:** Learning curves of the initial learning phase from trial 1 to 32. As the initial learning phase ended in 75% of the blocks before trial 18, estimates became increasingly unreliable after trial 17. **C:** Learning curves including trials of the initial learning phase and the subsequent practice phase. While the DRP, FOP and BP models became stationary when the initial learning phase ended, the Q-learning model further strengthened its associations between stimuli and responses, resulting in the best asymptotic fit on all three subsamples.

**Optimal and suboptimal errors.** While learning curves are typically used to characterize the temporal dynamics of learning processes, they are rather uninformative in terms of the circumstances by which different types of errors occurred during learning. To evaluate the generative performance of the models in terms of their ability to reproduce specific types of errors that occurred during initial learning, the errors were assigned to different categories. The first category consisted of ‘optimal errors’, defined as errors that occurred although the subject (or model) had chosen an optimal response, i.e. a response with maximal probability according to optimal (noise-free) FOP. The second category consisted of ‘suboptimal errors’, defined as errors that occurred for responses with nonzero, but not maximal probability according to noise-free FOP. Using these definitions, we found that the DRP models generated error distributions similar to those actually observed in the DRP subsample, whereas the FOP, BP and Q-learning models could not reproduce the actual distributions (see Figure 6). Specifically, the variability of the number of optimal errors generated by the FOP, BP and Q-learning models was much lower than actually observed. Moreover, these three models produced considerably more suboptimal errors than actually observed within the DRP and FOP sub-samples. The results of an extended analysis of error types, including errors that could have been avoided completely with optimal play, can be found in Supplementary Figure S3.

**Fig. 6.**
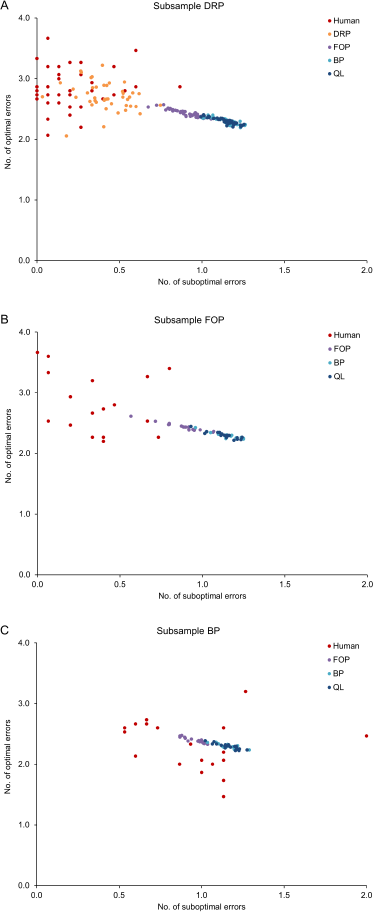
Joint distributions of optimal and suboptimal errors for the three subsamples. Optimal errors were defined as errors occurring when a response with maximum probability of being correct was selected and followed by negative feedback. Suboptimal errors were defined as errors occurring when a response with nonzero probability of being correct, but not maximum probability of being correct, was selected and followed by negative feedback. For each subject, the actual and modeled number of errors was averaged across learning blocks, i.e. each data point represents mean values of an individual subject. **A:** For the DRP subsample, the DRP models generated error distributions similar to those produced by the subjects, whereas the variability of optimal errors and the average number of suboptimal errors produced by the FOP, BP and Q-learning models were markedly different from the observed human data. **B:** Within the FOP subsample, the FOP, BP and Q-learning models again failed to reproduce the variability of optimal errors and the average number of suboptimal errors observed in the actual data. **C:** For the BP subsample, the three models generated approximately the same number of suboptimal errors as the subjects, but again failed to reproduce the variability of optimal errors.

Why did the FOP, BP and Q-learning models only poorly fit the optimal and suboptimal errors? The Q-learning model acquired stimulus-response associations independently foreach S-R pair, hence it could not distinguish between optimal and suboptimal errors, as the computation of response probabilities required inferences across S-R pairs. Moreover, the BP model could also not distinguish between optimal and suboptimal errors, as by definition differences between optimal and suboptimal response probabilities were removed before response selection. While the FOP model could distinguish between optimal and suboptimal errors, and indeed produced slightly better fits for these two error types than the BP and Q-learning models (see Supplementary Figure S3), the variability of optimal errors was still considerably reduced compared to the actual data, but also compared to the DRP models (see Figure 6 and Supplementary Figure S3). The reason for this reduced variability is that the number of optimal errors produced by FOP is independent of the stimulus sequences. More specifically, under noise-free FOP, the distribution of the number of optimal errors invariably converges towards the distribution shown in Supplementary Figure S4B for any stimulus sequence. In contrast, for the DRP models, the number of optimal errors varies as a function of the stimulus sequences, as shown in Supplementary Figure S4A.

## Discussion

Using computational models to analyze the initial learning phase of a trial-and-error learning task with deterministic feedback and hidden stimulus-response dependencies, we found that about 50% of the subjects employed deterministic response patterns to increase learning efficiency. Most of the remaining subjects either showed a tendency towards generic optimal learning, or performed better than predicted by pure associative learning by partially exploiting stimulus-response dependencies. A detailed analysis of specific error types showed that only the DRP model could generate the variability found in the human data, whereas the other three models were unable to reproduce this variability.

We followed a modeling approach that has been employed by a variety of studies before (2, 5, 8–10): The standard Q-learning model served as a baseline for comparison with more sophisticated models that either partially exploited task structure (BP) or approximated optimal performance (FOP), and found that the more sophisticated models provided a better fit to the data than the standard reinforcement learning model. This finding is in line with other studies that have compared pure associative learning with more sophisticated learning strategies in settings with probabilistic feedback (7, 14, 15, 22, 24), deterministic feedback (17, 21), or both types of feedback (28). Specifically, we found that the BP model provided a better fit to the data than the Q-learning model for a significant fraction of the subjects, which can be unambiguously attributed to certain inferences based on the one-to-one property of the stimulus-response mappings. More specifically, the BP model differed from Q-learning with respect to errors that could have been avoided by excluding responses that had been assigned to other stimuli in previous trials, as indicated by marked differences in specific error categories between these two models across all three subsamples (that is, error categories ‘correct for different stimulus’, ‘both correct for different stimulus and repeatedly wrong’ and ‘neither correct for different stimulus nor repeatedly wrong’, see Supplementary Figure S3). Hence, there is good evidence that these subjects exploited the fact that once a response had been assigned to a stimulus, it could be excluded for other stimuli. Similar findings have been reported before for trial-and-error learning tasks featuring two stimuli and probabilistic feedback (8–10, 14, 15, 22, 23).

More surprisingly perhaps, the BP model was outperformed by the FOP model on another significant fraction of the sample. This can be unambiguously attributed to differences in optimal and suboptimal errors, since the two models performed similarly with respect to other error types. These differences indicate that subjects of the FOP subsample did not only exclude responses previously assigned to other stimuli, as reflected by BP, but also exploited more subtle S-R-O dependencies corresponding to FOP; for instance the fact that when a response was rejected for some stimulus, its probability of being correct for one of the remaining open stimuli increased compared to the other available responses (cf. ‘After trial 1’ in Figure 2). Similar trends towards optimal task performance based on the integration of task structure into learning strategies have been reported before (3–5, 8).

The novel contribution of the results presented here is that they demonstrate that human learning strategies can be characterized beyond a general trend towards the optimal learning strategy. For 50% of the subjects, the initial learning phase was better explained by the DRP models than by FOP. Thus, these subjects did not select responses arbitrarily from the set of theoretically optimal responses, as predicted by FOP, but instead implemented a response selection procedure that determined a unique response in every trial. On the presented trial-and-error learning task with deterministic feedback, this was a highly adaptive learning strategy: Although being equivalent to FOP from a theoretical point of view, DRPs were more efficient from the human perspective as they considerably reduced working memory and computational demands. Indeed, using DRPs, only the correct or designated response for each stimulus had to be maintained in working memory, whereas FOP required tracking all 24 SR mappings. Computational costs were also significantly reduced, as the DRPs only required counting up to the next free response in case of negative feedback or storing the correct response in case of positive feedback, whereas FOP required computing response probabilities by averaging across all SR mappings consistent with the S-R-O history. Moreover, subjects could choose their preferred response order, which was arbitrary from a theoretical point of view, but not from the human perspective, as evidenced by the strongly nonuniform distribution across response orders (Supplementary Figure S1).

In order to successfully employ DRPs, subjects were required to reliably update their internal representations of task states on a trial-to-trial basis. Such an explicit and rapid updating of S-R-O contingencies, involving high-level cognitive processes and especially short-term maintenance of S-R-O information in working-memory, has also been reported before in studies on instruction-based and one-shot learning (5, 29–32). In these learning paradigms, subjects were either explicitly instructed on S-R contingencies (33–36), or had to infer instantaneously the correct response (37–39) or S-O causalities (5) in a single trial. Specifically, by investigating different types of S-R-O contingencies (40, 41) and learning conditions (42, 43), several studies have shown that the explicit instruction of S-R-O contingencies facilitates an almost error-free task performance right from the start of the practice phase. In the light of these studies, the findings presented here suggest that subjects employing DRPs might have divided the trial-and-error learning task implicitly into an exploration phase where they established the correct S-R links (equivalent to the initial learning phase defined for the computational models), and a subsequent practice phase where the S-R associations were consolidated via repetition. Together with the good asymptotic fit of the Q-learning model on the practice phase, our findings suggest that the high-level cognitive system supporting stimulus-response processing during initial learning is successively replaced by an associative system performing automatized, low-level stimulus-response transformations.

## Conclusion

Using a computational modeling approach, we showed that the subjects performed the presented trial-and-error learning task using highly adaptive and efficient learning strategies. While 50% of the subjects implemented deterministic response strategies in order to optimize task performance while keeping memory and computational demands bounded, most of the remaining subjects showed a general tendency to exploit hidden stimulus-response dependencies. These sophisticated learning strategies go beyond the incremental reinforcement of stimulus-response associations via feedback, and instead reflect the engagement of high-level cognitive processes during the initial learning phase.

## Acknowledgements

This work was funded by the German Research Foundation (Deutsche Forschungsgemeinschaft, DFG), SFB 940, subprojects Z2 and A2.

## Supplementary Figures

**Fig. S1.**
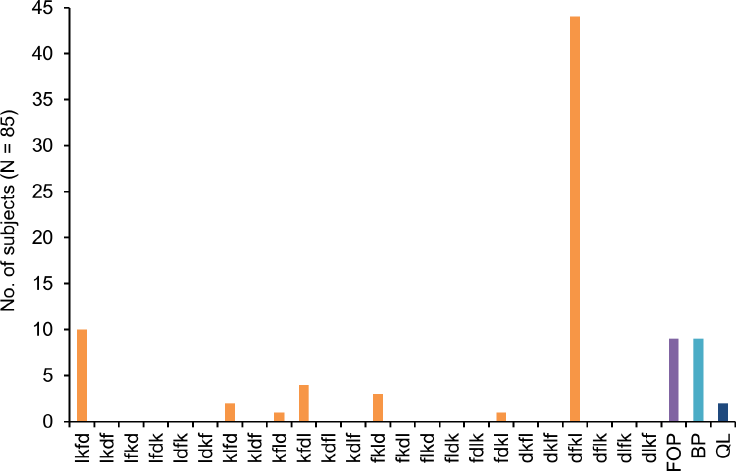
Preliminary model comparison including all 24 DRP response orders and the FOP, BP and Q-learning models. For each subject, it was determined which of the 27 models provided the largest log-likelihood score based on response data of the initial learning phase. Most subjects were best fitted either by the DRP dfkl, DRP lkfd, FOP or BP models. The response orders dfkl and lkfd correspond, respectively, to going from left to right and from right to left on the computer keyboard, which seem to be reasonable response strategies from a human perspective (while from a theoretical perspective, all 24 response orders are equivalent). In contrast, the third-ranked DRP response order kfdl corresponds to the rather implausible sequence right index finger, left index finger, left middle finger, right middle finger, and the fourth-ranked response order fkld corresponds also to an implausible sequence (left index finger, right index finger, right middle finger, left middle finger). Note that the preliminary model comparison reported here is based on the best-ranked model for each subject, with the difference between the best and second best model potentially being arbitrarily small. Thus, the fact that for a few subjects some implausible response orders obtained the highest log-likelihood score seems to reflect a bias in the model comparison procedure: Simply by submitting a larger number of models from the same class to the model comparison procedure, it becomes more likely that a model of this class obtains the highest score. To remove this bias from subsequent analyses, we constrained the model space to the five models DRP dfkl, DRP lkfd, FOP, BP and Q-learning, and conducted statistical tests for model comparison, reported in the main text.

**Fig. S2.**
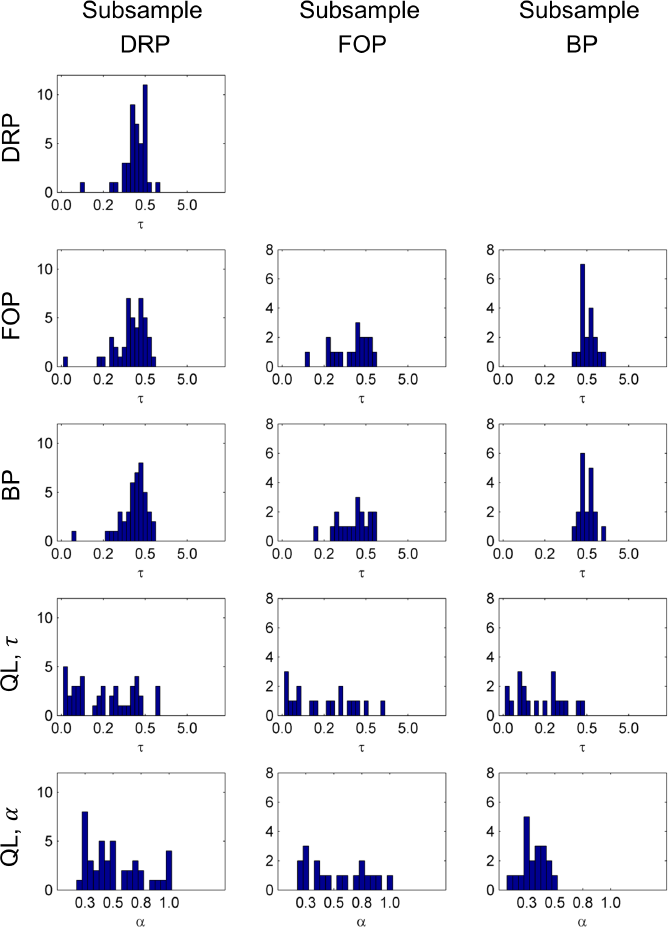
Maximum likelihood estimates of the model parameters, shown separately for the three subsamples. Response selection noise *τ* was fitted for all four models DRP, FOP, BP and Q-learning, while the learning rate *α* was only included in the Q-learning model. Response selection noise *τ* was optimized along the range 0,1/6,*1/5.8,…*, 1/0.2 (31 values), and the learning rate *α* was selected from the range 0.05,0.10,*…*, 1.0 (20 values). Parameters were fitted separately for each subject on response data of the initial learning phase.

**Fig. S3.**
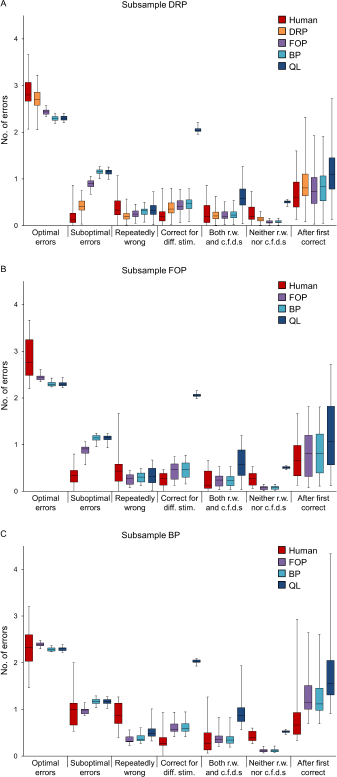
Extended analysis of error types of the initial learning phase. Errors were categorized into 7 different types. Optimal errors were defined as errors that occurred when a response with maximum probability of being correct was selected. Suboptimal errors were defined as errors that occurred when a response with nonzero probability, but not maximal probability, was selected. Errors were categorized as ‘repeatedly wrong’ if negative feedback had been received before for the respective S-R pair. Errors were categorized as ‘correct for a different stimulus’ if a response was selected that had been assigned to a different stimulus in earlier trials. Errors were categorized as ‘both repeatedly wrong and correct for a different stimulus’ if both criteria were fulfilled. Errors were categorized as ‘neither repeatedly wrong nor correct for a different stimulus’ if an indirect inference would have led to the correct response, as for example in step 5 of Figure 3, where the correct response for the fourth stimulus was inferred based on the one-to-one property of the S-R mappings. Finally, errors were categorized as ‘after first correct’ if the respective S-R pair had been performed correctly before. For each subject, the number of errors of each type was averaged across the 15 learning blocks of interest. The plots show median, first- and third quartile, and minimum and maximum values across the subjects of the respective sample. **A:** Data of the DRP subsample. As also depicted in Figure 6 of the main text, the DRP models performed considerably better than the other models in terms of optimal and suboptimal errors. The FOP model showed at least a tendency towards the actual data for these two error types, but all three models (FOP, BP and Q-learning) failed to reproduce the high variability of optimal and suboptimal errors found in the actual data. Moreover, the Q-learning model was unable to exploit the one-to-one property of the S-R mappings, as can be seen by the high rate of ‘correct for a different stimulus’, ‘both repeatedly wrong and correct for a different stimulus’ and ‘neither repeatedly wrong nor correct for a different stimulus’ errors. **B:** Data of the FOP subsample. As in A, the FOP model showed a slightly better fit in terms of optimal and suboptimal errors than the BP and Q-learning models, but none of the three models could reproduce the high variability of these error types. **C:** Data of the BP subsample. Again, all three models showed much lower variability in terms of optimal and suboptimal errors than observed in the actual data.

**Fig. S4.**
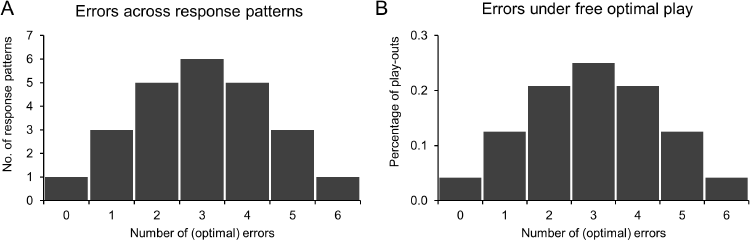
Distributions of optimal errors under optimal (noise-free) play. **A:** For any stimulus sequence, the number of optimal errors produced by the 24 response orders invariably resulted in the shown distribution. **B:** For a large number of repetitions, the number of errors under free optimal play converged to the same distribution as in A on any stimulus sequence.

## Appendix

**Notation.** Experimental trials are indexed with *t* = 1,2,3,…, and each trial consists of a stimulus *s_t_* ∈ {*1,2,3,4*}, a response *r_t_* ∈ {*d, f, k, l*} and an outcome *o_t_* ∈ {*0,1*}. After completion of trial no. t, the history of stimuli, responses and outcomes is denoted as *H_t_* = {(*s*)*_t_*, (*r*)*_t_*, (*o*)*_t_*}, with (*s*)*_t_* = (*s*_1_,*s*_2_,…,*s_t_*), (*r*)*t* = (*r*_1_,*r*_2_,…,*r_t_*), (*o*)*_t_* = (*o*_1_,*o*_2_,…,*o_t_*).

Let {*M_i_*|*i* = 1,2,3,…,24} be the set of the 24 one-to-one mappings between four stimuli and four responses as shown in Figure 2 in the main text, with deterministic outcomes being either zero or one. In a probabilistic framework, the one-to-one property of S-R mappings with deterministic outcomes translates into: For a given mapping *M_i_*, for each stimulus *ŝ*, there exists a response 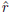 such that 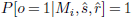 and 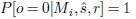 for all responses 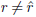 and 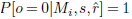 for all stimuli *s* ≠ *ŝ*.

We will show that in order to maximize expected reward and minimize expected uncertainty, it is sufficient to track the probability distribution over the S-R mappings *M_i_, i* = 1,2,…, 24 given the history of S-R-O combinations, that is, it is sufficient to track *P* [*M_i_\Ht*] for all *i* = 1,2,…, 24. Initially, without any S-R-O information available, we assume that all mappings are equally likely, that is, we define *P* [*Mi\H o*] = 1 /24 for all *i* = 1,2,…, 24.

**Computation of outcome probabilities.** First, we show that when stimulus s*_t_* is presented, the likelihood for getting an outcome *o*_t_ by responding with *r*_t_ can be computed as:

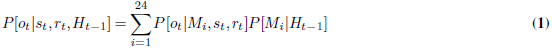

It is:

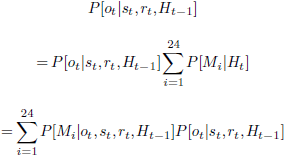

As generally *P* [*A | B, C*]*P* [*B | C*] = *P* [*B | A,C*]*P* [*A | C*], it follows with *A* = *M_i_*, *B* = *o_t_* and *C* = {*s_t_, r_t_, H_t_*–*i*} that

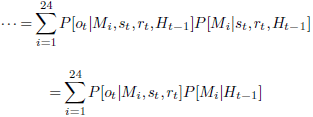

**Maximization of expected reward.** As *P*[*o_t_ |M_i_,s_t_,r_t_*] is known by the definition of *M_i_*, it is sufficient to have *P*[*M_i_*|*H_t_*_–*i*_] at hand in order to maximize expected reward in trial *t:*

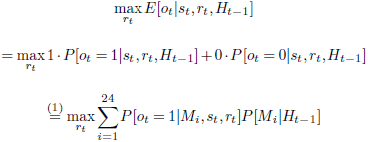

**Updating.** After having obtained an outcome *o_t_* for response *r_t_*, the likelihood for each mapping *M_i_* can be updated in the following way:

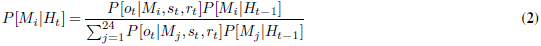

From this identity, we can see that by starting with a uniform distribution, the distribution over the *M_i_*’s (i.e. *P*[*M. |H_t_*]) evolves as a mixture of a uniform distribution over a subset of *M_i_*’s and zeros for the rest of the *Mi*’s, that is, after trial *t* exists an *n_t_* ∈ {1,2,…, 24} and subindexes {*i_1_,i_2_,…,i_nt_*}, such that *P* [*M_i_ | H_t_*] = *1/n_t_* for all *i* ∈ {*i_1_,i_2_,…,i_nt_*} and *P* [*M_i_ | H_t_*] = 0 for all *i* ∈ {1,2,…, 24},*i*_2_,…,*i_nt_*}. To this end, we assume the properties hold for the previous trial, i.e. it is *P*[*Mi|Ht-i*] ∈ {*0,1/nt-i*} for all i = 1,2,…,24. Generally, it is *P*[*ot|Mi,st*,*n*] ∈ {0,1}. If follows directly from (2) that *P*[*Mi|Ht*] =0 if *P*[*Mi|Ht-i*] =0 or *P*[*ot|M_h_st*,*n*] = 0. Hence, we assume that *P*[*Mi|Ht-i*] = 1/*nt-i* and *P*[*ot|Mi,st,rt*] = 1. We define *nt* = #{*j* ∈ {1,2,…,24}|*P*[*ot|Mj,s_u_n*] = 1,*P*[*Mj|Ht-i*] = 1/*nt-i*}., Then, using (2) again, it is 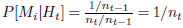. Thus, for all *i* ∈ {*1,2,…, 24*}, it is *P* [*Mi | Ht*] ∈ {*0,1/nt*}.

To equation (2): Generally, it is

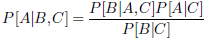

With *A = M_i_, B = o_t_* and *C* = {*s_t_,r_t_*, *H_t_*_–*i*_} it follows that

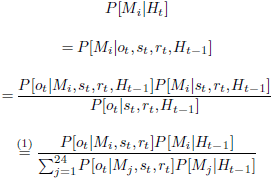

**Minimization of expected uncertainty.** Next, we show that maximizing expected reward in the current trial also minimizes the expected uncertainty. To this end, at the beginning of trial t, we define the set of S-R mappings that are consistent with the S-R-0 history: *c_t–i_* = {*i* ∈{1,2,…,24}|*P*[*M_i_|H_t-i_*] > 0} and *n_t-i_* = ≠*c_t-i_*. Moreover, given the currently presented stimulus s_t_, we define for the four responses d, f, k, l the sets of S-R mappings that are consistent with a positive outcome:

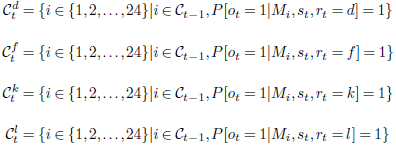

And accordingly, we define 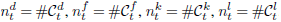.

From equation (1) follows, with response *x ŝ {d, f, k, l}:*

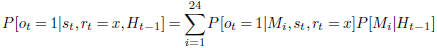

Moreover, as we have seen above, S-R mappings in *C_t-i_* are equally likely given *H_t_*_–*i*_ (i.e. *P*[*M_i_|H_t–i_*] = 1/*n_t–i_* for all *i* ∈ *C_t–i_*), hence we get for *x* ∈ {*d, f, k, l}:*

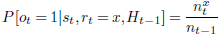

Additionally, from the one-to-one property of the S-R mappings follows that

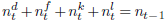

Importantly, the updating procedure from equation (2) implies that when response *x* (with *x ∈ {d, f, k, l*}) is given (i.e. *rt = x*) and followed by a positive outcome (i.e. *ot =* 1), then 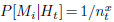 for 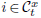 and *P*[*M*_*i*_|*H*_*t*_| = 0 for 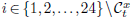, whereas if the response is not rewarded (o_t_ *=* 0), then 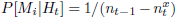 for 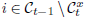 and *P*[*M_i_|H_t_] = 0* for 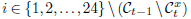.

Without loss of generality, we assume that response *d* maximizes expected reward:

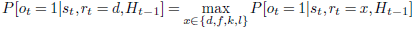

Equivalently, it is *n_d_* = *max{n_d_,n_f_,n_k_,n_l_*}. We want to show that response *d* also minimizes the expected uncertainty of the S-R mappings, with uncertainty being defined as the entropy *H* of the distribution *P*[*M. |H_t_]*, i.e. we want to show:

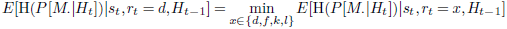

Given the considerations above, it follows that:

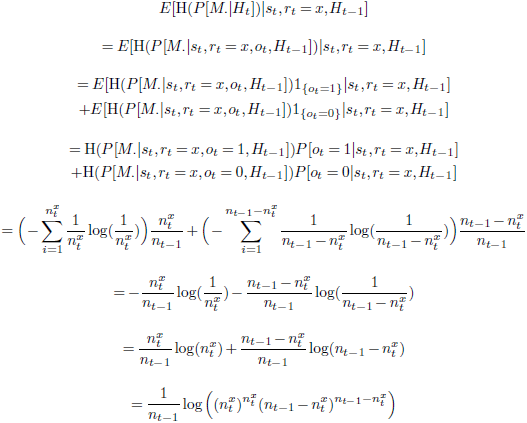

We show exemplarily for response *f* that from *n_d_* ≥ *n_f_* follows that

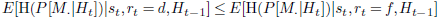

Exploiting the transformation from above, we have to show that

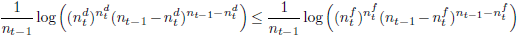

Which can be simplified further as follows:

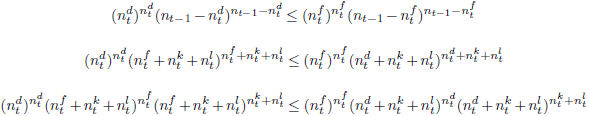

If 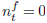 the inequality simplifies to

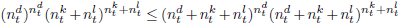

which is correct, given that 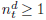 and 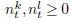.

Hence, we continue with 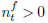:

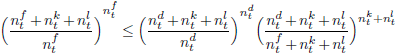

From *n_d_* > *nf* follows that

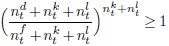

Hence, it remains to show that

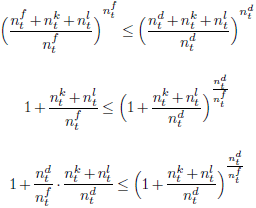

With 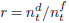 and 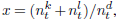 this is equivalent to Bernoulli’s inequality:

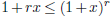

**Maximization of expected information gain.** Finally, a response that maximizes the expected reward also maximizes the expected information gain, with information gain being defined as the Kullback-Leibler divergence between *P [M. | H_t_*] and *P*[*M. |H_t–i_*]. As above, we assume without loss of generality that *n_d_* = *max*{*n_d_,nf,n_k_,n_l_*}, and exemplarily show for response *f* that:

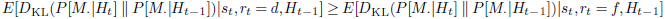

Generally, for response *x* ∈ {*d, f, k, l*}, it is:

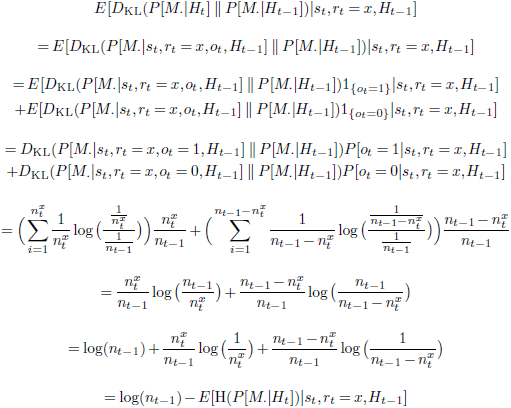

Hence, in order to show that

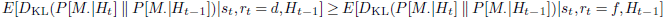

it is sufficient to show that

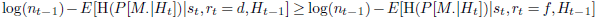

or equivalently, that

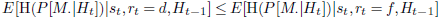

which we have already shown above.

